# Netpredictor: R and Shiny package to perform Drug-Target Bipartite network analysis and prediction of missing links

**DOI:** 10.1101/080036

**Authors:** Abhik Seal, David J. Wild

## Abstract

Netpredictor is an R package for prediction of missing links in any given bipartite network. The package provides utilities to compute missing links in a bipartite and well as unipartite networks using Random Walk with Restart and Network inference algorithm. The package also allows computation of Bipartite network properties, visualization of communities for two different sets of nodes, and calculation of significant interactions between two sets of nodes using permutation based testing. The R standalone package (including detailed introductory vignettes) and associated R Shiny web application is available under the GPL-2 Open Source license and is freely available to download from github Netpredictor repository and Shiny Netpredictor repository respectively.

## Introduction

Identifying missing associations between drugs and targets provides insights into polypharmacology and off-target mediated effects of chemical compounds in biological systems. Traditional machine learning algorithms like Naive Bayes, SVM and Random Forest have been successfully applied to predict drug target relations [1–4]. However, using supervised machine learning methods requires training sets, and they can suffer from accuracy problems through insufficient sampling or scope of training sets. During the last years, the field of semi-supervised learning has been applied to methods based on graphs or networks. The data points are represented as vertices of a network, while the links between the vertices depend upon the labeled information. Thus, it is desirable to develop a predictive model based on using both labeled and unlabeled information. Network based models can avoid these kind of issues. The main advantages of network-based methods are:

- They use label information and as well as unlabeled data as input in the form of vectors.
- Once can use multiple classes inside the network structure.
- It uses multitude of paths to compute associations.
- Network based methods mostly use transductive learning strategy,in which the test set is unlabelled but while computation it uses the information from neighbourhood.

**Table 1.**
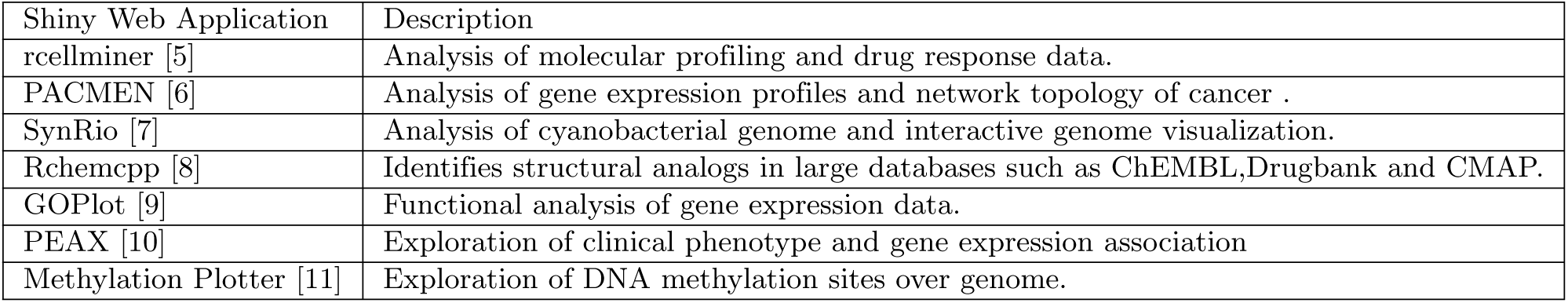
Table shows some lifescience related applications developed in R and shiny

With the advent of the R open source statistical programming language [12] and the gaining popularity of the RShiny package for interface development around R [13] it has become straightforward for programmers to create and deploy web applications on windows and Linux servers. R and RShiny have already used in several biomedical applications. Table 1 shows some of these. We used R and R shiny to create Netpredictor standalone and web application respectively, which is freely available and open source. The web application framework in R allows creation of a simple intuitive user interface with dynamic filters and real-time exploratory analysis. Shiny also allows integration of additional R packages, Javascript libraries and CSS for customization. Web applications are accessible via browser or can be run locally on the user’s computer. The R package described in this paper provides utilities to compute recommendations in a bipartite network and well as unipartite network based on HeatS [14], Random walk with Restart (RWR), Network based inference (NBI) [15] and combination of RWR and NBI(netcombo). In order to understand the topology of the network, the package also provides ways to compute bipartite network properties such as degree centrality, density of the network, betweenness centrality, number of sets of nodes and total number of interactions for given bipartite network. The package also performs graph partitioning such as bipartite community detection using the lpbrim algorithm [16, 17] and visualization of communities, network permutations to compute the significance of predictions and performance of the algorithms based on user given data.

We implemented two network algorithms in this package NBI and RWR. Cheng [18] developed a technique based on Network Based Inference (NBI) and developed three supervised methods on drug similarity, target similarity and network topology and showed superior performance of network topology based method. Alaimo [19] extended Cheng’s method to integrate chemical and target similarity and showed that that the performance of the method is superior to Cheng’s model. Chen [20] and Seal [21] have used random walk with restart (RWR) based method to predict drug target interactions on a heterogeneous network made up of drug-drug similarity, protein-protein similarity and bipartite graph between drugs and targets. Seal et al. have extended the method by optimizing a parameter *η* which showed that the performance of RWR is independent of the choice of using Chemical fingerprint features.

## Design and Implementation

The Netpredictor package can be used in two ways - either in standalone form and compelling web application running locally or on an Amazon cloud server [22]. The web applications accessible through the Internet and standalone package are functionally identical. More details regarding the package accessibility and the instructions on how to use it via the web application and run locally are given in the availability section. The interface consists of two parts - a web interface and a web server. Both of these components are controlled by code that is written within the framework of Shiny application in R. RShiny uses “reactive programming” which ensures that changes in inputs are immediately reflected in outputs, making it possible to build a highly interactive tool. Within the RShiny package, ordinary controllers or widgets are provided for ease of use for application programmers. Many of the procedures like uploading files, refreshing the page, drawing new plots and tables are provided automatically. The communication between the client and server is done over the normal TCP connection. The data traffic that is needed for many of web applications between the browser and the server is facilitated over the websockets protocol. This protocol operates separately using handshake mechanism between the client and server is done over the HTTP protocol. The duplex connection is open all the time and therefore authentication is not needed when exchange is done. In order for an RShiny app to execute, we have to create an RShiny server. RShiny follows a pre-defined way to write R scripts. It consists of server.R and ui.R, which need to be in same directory location. If a developer wants to customize the user interface shiny can also integrate additional CSS and Javascript libraries within the web application. The GUI consists of introduction page with tab panels shown in (Fig 1 S2 Fig). The first tab, start prediction, consists of sidebar panels and a main output panel (Fig 2 S3 Fig). The sidebar is used to upload the data and select the algorithms and its parameters. The start prediction tab consists of data upload, compute recommendations, compute network properties and visualization of user given data. The advanced analysis tab has two sections the statistical analysis section and permutation testing tab. We also computed the recommendations of the Drugbank database using NBI and included the predictions results in the Drugbank search tab.

**Figure 1.**
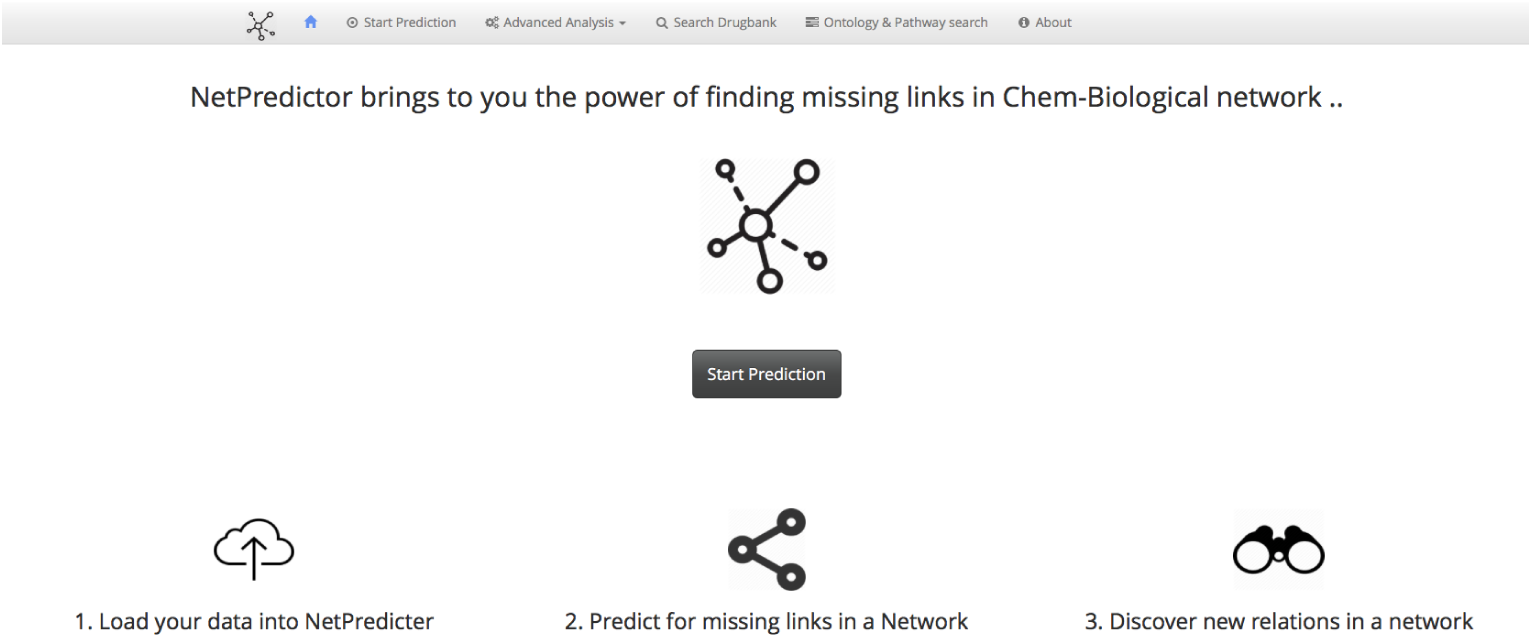
Figure shows the first page of the netpredictor tool build using R shiny.

## Main Features of netpredictor standalone and web tool

The standalone R package application can perform prediction on unipartite networks using a set of different similarity measures between vertices of a graph in order to predict unknown edges (links) [23, 24]. The prediction methods are classified into two categories:

- Neighborhood based metrics and
- Path based metrics

**Figure 2.**
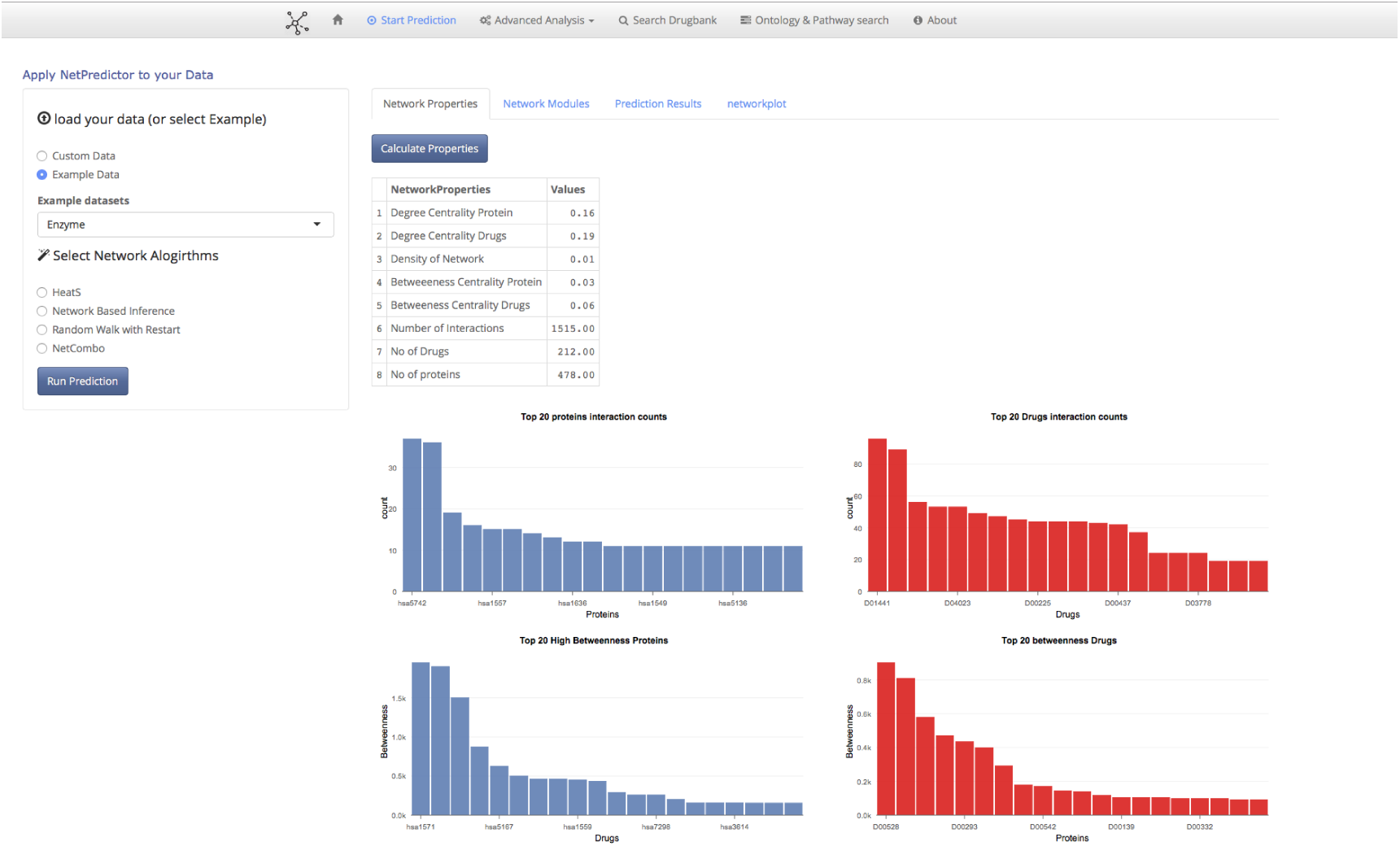
Figure Shows the Network properties tab.

For neighbourhood based metrics the methods which are implemented are (i) common neighbours (ii) jaccard coefficient [26] (iii) cosine similarity (iv) hub promoted index [27] (v) hub depressed index (vi) Adamic Adar index [28] (vii) Preferential attachment [29] Resource allocation [30] (ix) Leicht-Holme-Nerman Index [31]. Similarly using path-based metrics one can compute paths between two nodes as similarity between node pairs. The methods are:

i. The local path based metric [32] uses the path of length 2 and length 3. The metric uses the information of the nearest neighbours and it also uses the information from the nodes within length of 3 distances from the current node.
ii. The Katz metric [33] is based on similarity of all the paths in a graph.This method counts all the paths between given pair of nodes with shorter paths counting more heavily. Parameters are exponential.
iii. Geodesic similarity metric calculates similarity score for vertices based on the shortest paths between two given vertices.
iv. Hitting time [34] is calculated based on a random walk starts at a node x and iteratively moves to a neighbor of x chosen uniformly at random. The Hitting time H_*x,y*_ from x to y is the expected number of steps required for a random walk starting at x to reach y.
v. Random walk with restart [15, 34, 35] is based on pagerank algorithm [36]. To compute proximity score between two vertexes we start a random walker at each time step with the probability 1 - c, the walker walks to one of the neighbors and with probability c, the walker goes back to start node. After many time steps the probability of finding the random walker at a node converges to the steady-state probability.

The significance of interaction of links is based on random permutation testing. A random permutation test compares the value of the test statistic predicted data value to the distribution of test statistics when the data are permuted. Supporting Information S1 NetpredictorVignette provides tutorial for this netpredictor standalone R package. In the web application app one can load their own data or can use the given sample datasets used in the software. For the custom dataset option one needs to upload bipartite adjacency matrix along with the drug similarity matrix and protein sequence matrix. From the given datasets Enzyme, GPCR, Ion Channel and Nuclear Receptor in the application one can load the data and set the parameters for the given algorithms and start computations. The data structure the web application accepts matrix format files for computation.

A summary of the contents of each of the tabs shiny netpredictor application is reported in Table 2.

### Start Prediction Tab

The *start prediction* tab is designed to upload a network in matrix format and compute it properties, searching for modules, fast prediction of missing interactions, visualization of bipartite modules and predicted network. For the custom dataset, in the input drug-target binary matrix, target nodes should be in rows and drug nodes in the columns. The drug similarity matrix and the target similarity should have the exact number of drugs and targets from the binary matrix. For HeatS, only the bipartite network is used to compute the recommendation of links. For RWR, NBI, and Netcombo all of these require three matrices. The default parameters are already being set for the algorithms. The main panel of the start prediction tab has four tabs that compute network properties, network modules, the prediction results and predicted network plot.

Bipartite network properties are calculated by transforming the network in to one-mode networks (contain one set of nodes) called projection of the network in which a bipartite network of drugs and proteins two drugs are connected if they share a single protein similarly two proteins are connected if they share a single drug molecule. Using the two-projected network of drugs and proteins we compute degree centrality, betweenness, total number of interactions, total number of each of the nodes and distribution of the drug and target nodes shown in Figure 2. WE have implemented the visualization of cousts and betweenness histograms using the rCharts R package [37]. Bipartite network modules are computed using the lpbrim algorithm [38] for which lpbrim R package is used [17]. The algorithm consists of two stages. First, during the LP phase, neighboring nodes (i.e. those which share links) exchange their labels representing the community they belong to, with each node receiving the most common label amongst its neighbors. The process is iterated until densely connected groups of nodes reach a consensus of what is the most representative label, as indicated by the fact that the modularity is not increased by additional exchanges. Second, the BRIM algorithm (2) refines the partitions found with label propagation. HeatS and network based inference compute (NBI) recommendations using a bipartite graph, where a two phase resource transfer Information from set of nodes in A gets distributed to B set of nodes and then again goes back to resource A. This process allows us to define a technique for the calculation of the weight matrix W. HeatS uses only the drug target bipartite data matrix and NBI uses similarity matrices of drug chemical similarity matrix and protein similarity matrix. The random walk with restart(RWR) algorithm uses all the three different matrices to compute the recommendations. Netcombo computes both NBI and RWR and then averages the scores. The prediction results tab shows the computed results using the javaScript library DataTables [39]. The data table provides columns filters and search options. The network plot tab represent the network using the visNetwork R package [40] The Network visualization is made using vis.js javascript library. Javascript libraries can be integrated using a binding between R and javascript data visualization libraries (Fig 3 S4 Fig).

The htmlwidgets library [41] can generate a web based plot by just calling a function that looks like any other R plotting function.

**Table 2.** Table shows the functions of tabs in Shiny web application

| Interface Tabs | Description |
| --- | --- |
| Load data and select algorithms | The load data and selection of algorithms panel allows users to load custom data or example datasets in matrix format. Users need to upload the matrices binary drug-target bipartite network, drug –drug similarity and protein – protein similarity along with algorithm and parameters of choice. |
| Network Properties | The network properties several different properties of the bipartite graph such as the degree centrality of two types of nodes, density of the network, betweenness of two types of nodes, total number of interactions, count of each type of nodes. |
| Network Modules | Bipartite network modules are computed using the lpbrim algorithm [38] the tab shows the bipartite nodes as tables and module network. Modules are dynamically updated based on input data. |
| Prediction Results | For a given dataset to compute the results one can select any one of the algorithms . The results are shown using the jquery Data Tables library. The table shows the drugs , targets, pvalues, outcome ( True/predicted interaction). |
| Network Plot | The network plot tab plots the computed predicted network. It uses visNetwork package which uses the vis.js library to generate network. The predicted interactions are marked in dashed lines and true interactions are marked with bold lines. Drop downs are provided to select specific nodes and groups. |
| Statistical Testing | The statistical testing tab tests the performance of the model based one the random removal of true links from the network based on the frequency of the drug-target associations. It measures the auac, auc, auctop(10%), bedroc and enrichment of links. Based on these scores we select which algorithm to use. |
| Permutation Testing | In permutation testing significance of the associations are calculated by, randomly permuting the matrices and and compute the significance using, standard normal distribution. |
| Search Drugbank | This tab allows users to search predicted drug target associations from the drugbank database from 5970 drugs and 3797 proteins from a total of 316645 predicted and 14167 true interactions. |
| Ontology and Pathway Search | This tab allows users to search for enrichment of Ontologies and Pathways using a given set of genes. |

**Figure 3.**
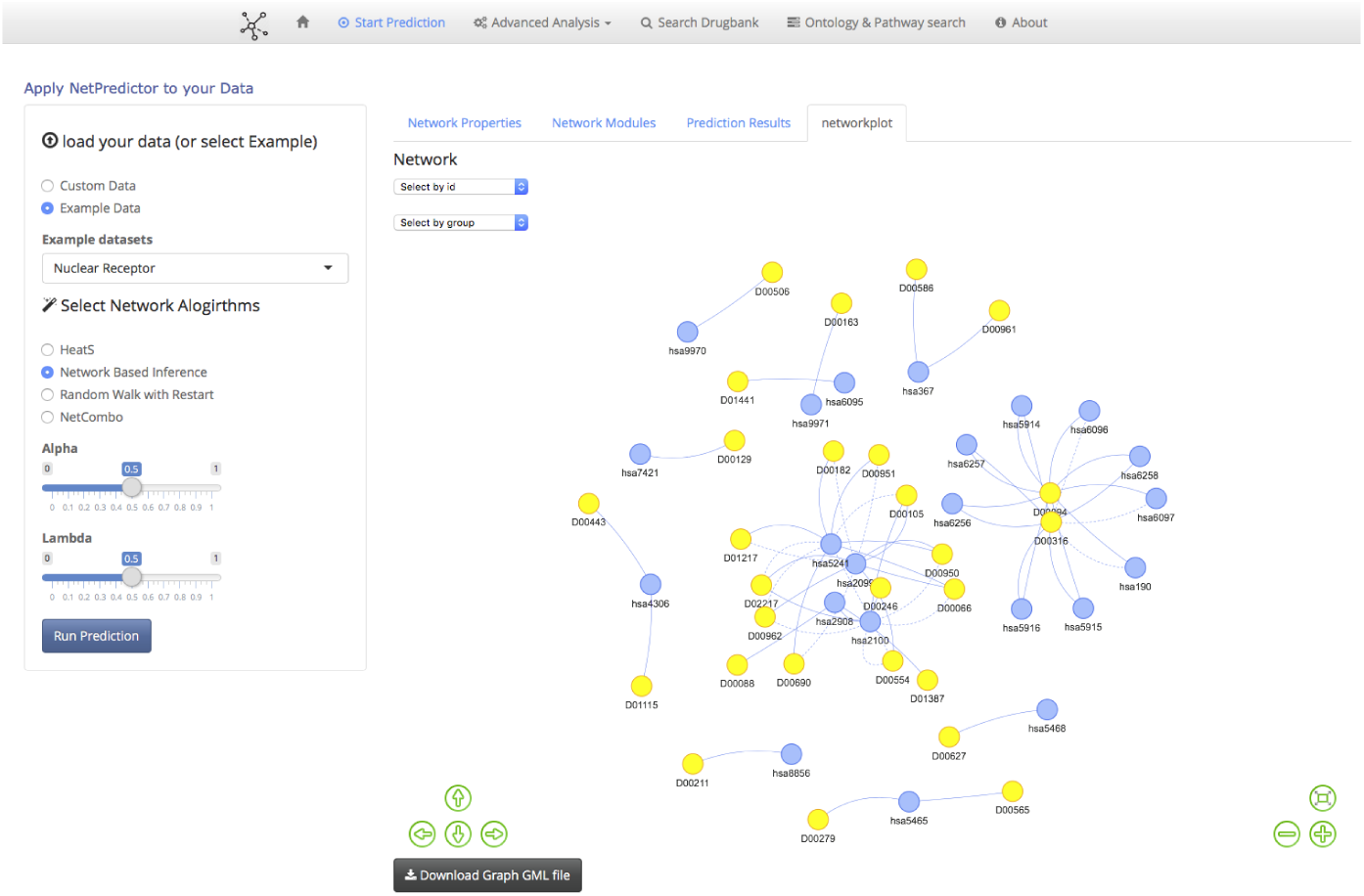
Figure shows predicted network plot.

### Advanced Analysis tab

The *advanced analysis* consist of two functions the statistical analysis tab and permutation testing. The statistical analysis tab computes the performance of the algorithms. Three algorithms are network based inference, random walk with restart and netcombo can be used. One can randomly remove the true links from the network using frequency of the drug target interactions in the network. The performance of the algorithm is checked when the removed links are repredicted. The statistics used to evaluate the performance is AUAC, AUC, AUCTOP(10%), Boltzmann-enhanced discrimination of ROC (BEDROC) [42] and enrichment factor(EF). The data table gets automatically updated for each of the computations. The results are reported in main panel using data tables. The significance of interactions using random permutations can be computed for the given network using network based inference and random walk with restart. The networks are randomized and significance of the interactions are calculated based on standard normal distribution. The user needs to give total number of permutations to compute and the significant interactions to keep.

### Search Drugbank tab

The *drugbank tab* (Fig 4 S5 Fig) helps to search predicted interactions computed using NBI method using the drugbank database [43]. One can search for targets given a specific drugbank ID and search for drugs given a specific hugo gene name [44].In fig3 the data table shows the drug target significant scores whether it is a true or predicted interaction, Mesh categories of drugs, ATC Codes and groups (approved, illicit,withdrawn, investigational, experimental). Currently the drugbank search tab only supports data computed using Netowrk based inference. The computed results and the associated meta-data are stored in a sqllite database [45] for access through shiny data tables interface.

**Figure 4.**
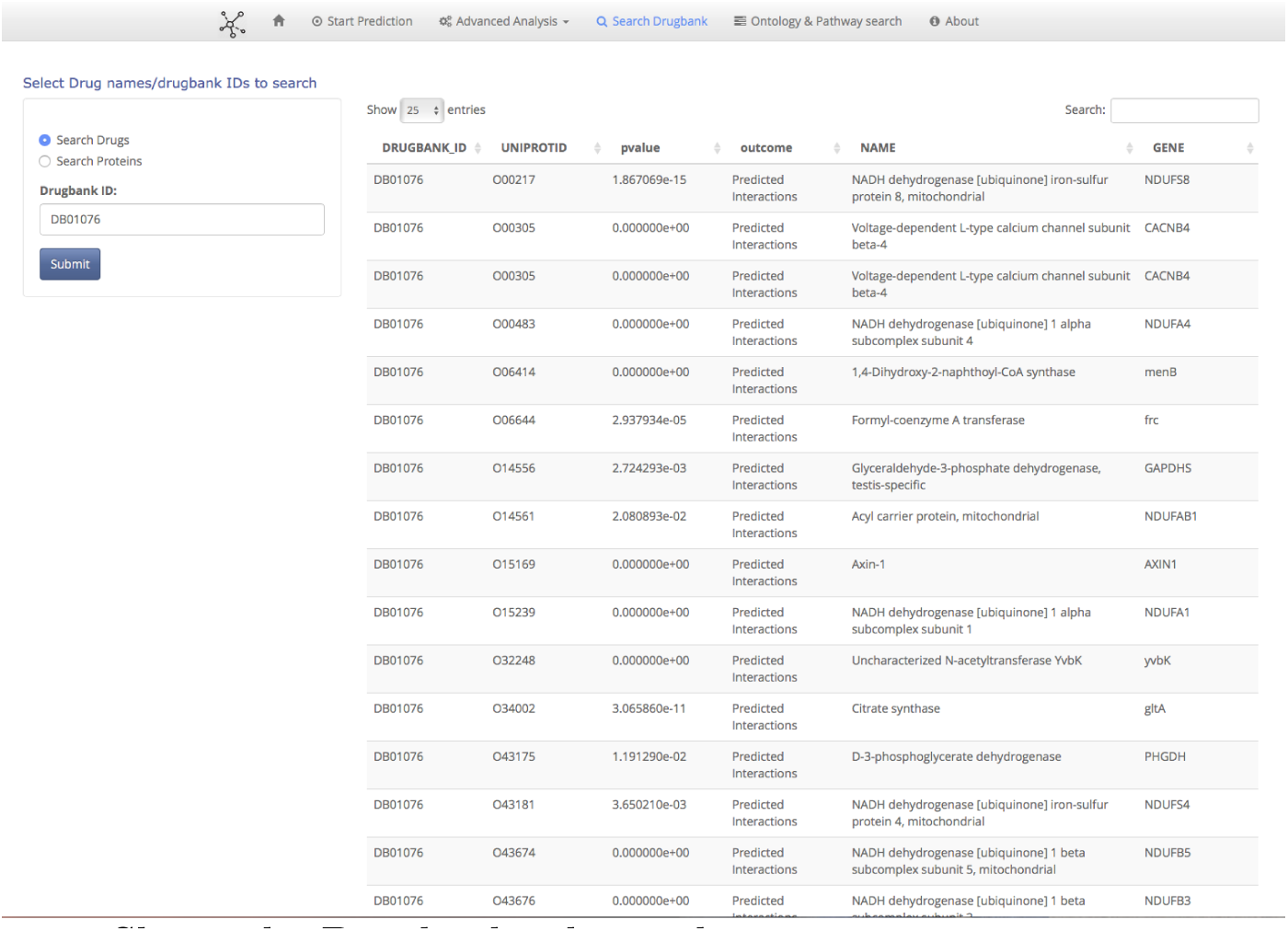
Shows the Drugbank tab panel.

### Ontology and Pathway search tab

The *Ontology and pathway search tab*(Fig 5 S6 Fig) helps to search the relevant gene ontology terms and pathways for a given set of genes. A search can be made based on predicted proteins and in order to understand its function, location and pathway this tab can help to understand it. The level of ontology can also be given to the user input. We used biomart services using the biomaRT R package [46] to convert genes names to entrez ids and then the clusterProfiler R package [47] to retrieve the gene ontology lists. The pathway enrichment is based on the ReactomePA R package [48].

**Figure 5.**
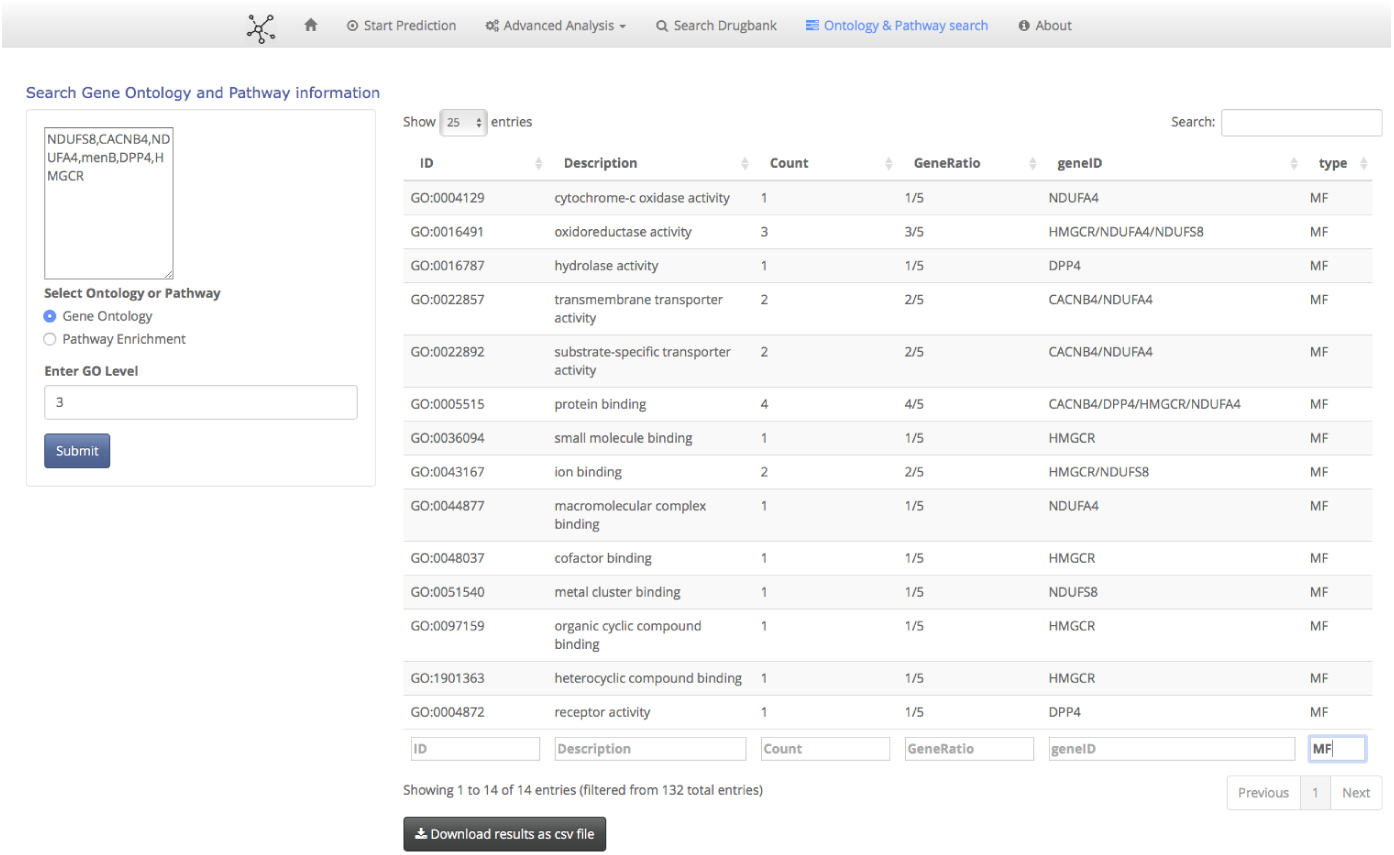
Shows the Ontology and Pathway search tab panel.

**Table 3.** Table shows the performance of RWR and NBI on different datasets

| AUAC | AUC | AUCTOP | BDR | EFC | Dataset | Method |
| --- | --- | --- | --- | --- | --- | --- |
| 0.934 | 0.899 | 0.577 | 0.506 | 8.38 | Enzyme | RWR |
| 0.823 | 0.882 | 0.274 | 0.252 | 5.066 | GPCR | RWR |
| 0.841 | 0.88 | 0.283 | 0.254 | 6.28 | Ion Channel | RWR |
| 0.585 | 0.698 | 0.018 | 0.089 | 0.295 | Nuclear Receptor | RWR |
| 0.834 | 0.879 | 0.636 | 0.429 | 4.633 | Enzyme | NBI |
| 0.767 | 0.466 | 0.281 | 0.217 | 4.3 | GPCR | NBI |
| 0.874 | 0.938 | 0.349 | 0.284 | 6.756 | Ion Channel | NBI |
| 0.487 | 0.52 | 0.049 | 0.05 | 0.309 | Nuclear Receptor | NBI |

## Results

In this section we illustrate the use of Netpredictor package in prediction of drug target interactions and analysis of networks. The information about the interactions between drugs and target proteins was obtained from Yamanishi etal [49] where the number of drugs 212, 99, 105 and 27, interacting with enzymes, ion channels, GPCRs and nuclear receptors respectively. The numbers of the corresponding target proteins in these classes are 478, 146, 84 and 22 respectively. The numbers of the corresponding interactions are 1515, 776, 314 and 44. We performed both network based inference and Random walk with restart on all of these datasets. To check the performance we randomly removed 20% of the interactions from each of the dataset and computed the performance 50 times and calculated the mean performance of each of these methods. The results are given in Table 3. Clearly, RWR supersedes its performance compared to network based inference in Enzyme and the GPCR dataset. However, computation of NBI algorithm takes less amount of time than RWR. For the drugbank tab we download the latest drugbank set version 4.3 and created a drug target interaction list of 5970 drugs and 3797 proteins We computed similarities of drugs using RDkit [51] ECFP6 fingerprint and local sequence similarity of proteins using smith waterman algorithm and normalized using the procedure proposed by Bleakley and Yamanishi [52] and integrated the matrices for network based inference computation. We ran the computations 50 times and kept the significant drug target relations(p ≤ 0.05) where a total of 316645 predicted interactions and 14167 true interactions present in the system.

## Availability and Future Directions

In this paper we presented netpredictor, a standalone and web application for drug target interaction prediction. Netpredictor uses a shiny framework to develop web pages and the application can be accessed from web browsers. To set up the Netpredictor application locally there are some additional requirements other than shiny which are given below,

- Firstly, the user has to have the R statistical environment installed, for which instructions can be found in R software home page.
- Secondly, the devtools R package [50] has to be installed. The package can be installed using `install. packages` (“devtools”)
- Also for fast computation Microsoft R Open package needs to be installed which can be obtained from https://mran.revolutionanalytics.com/documents/rro/installation/. Microsoft R Open includes multi-threaded math libraries to improve the performance of R. R is usually single threaded but if its linked to the multi-threaded BLAS/LAPACK libraries it can perform in multi-threaded manner. This usually helps in matrix multiplications, decompositions and higher level matrix operations to run in parallel and minimize computation times.
- After installing R, R open and shiny calling `shiny::runGitHub (‘Shiny NetPredictor’, ‘abhik1368’)`

This will load all the libraries need to run netpredictor in browser. The application can be accessed in any of the default web browsers. The netpredictor R package (https://github.com/abhik1368/netpredictor) and the Shiny Web application(https://github.com/abhik1368/Shiny_NetPredictor) is freely available.

Users can follow the “Issues” link on the GitHub site to report bugs or suggest enhancements. In future the intention is to include Open Biomedical Ontologies for proteins to perform enrichment analysis. The package is scalable for further development integrating more algorithms.

## Author contributions

Conceived and designed the experiments and tool: AS. Performed the experiments: AS Analyzed the data: AS. Contributed reagents/materials/analysis tools: AS,DJW. Wrote the paper: AS,DJW. Interpreted the results, drafted the manuscript and contributed to revisions: AS,DJW. Read and approved the final manuscript: AS,DJW.

## References

1. Cao DS, Liang YZ, Yan J, Tan GS, Xu QS, Liu S (2013b). “PyDPI: Freely Available Python Package for Chemoinformatics, Bioinformatics, and Chemogenomics Studies.” Journal of chemical information and modeling.

2. Cao DS, Liang YZ, Deng Z, Hu QN, He M, Xu QS, Zhou GH, Zhang LX, Deng Zx, Liu S (2013a). “Genome-Scale Screening of Drug-Target Associations Relevant to Ki Using a Chemogenomics Approach.” PloS one, 8(4), e57680.

3. Proteochemometric modeling as a tool to design selective compounds and for extrapolating to novel targets GJP van Westen, JK Wegner, AP IJzerman, HWT van Vlijmen, A Bender MedChemComm 2 (1), 16–30

4. Proteochemometric modelling coupled to in silico target prediction: an integrated approach for the simultaneous prediction of polypharmacology and binding affinity/potency of small molecules. S Paricharak, I Cortés-Ciriano, AP IJzerman, TE Malliavin, A Bender J. Cheminformatics 7, 15

5. Luna A, Rajapakse VN, Sousa FG, Gao J, Schultz N, Varma S, Reinhold W, Sander C, Pommier Y. rcellminer: exploring molecular profiles and drug response of the NCI-60 cell lines in R. Bioinformatics. 2015 Dec 3. pii: btv701

6. Ghazanfar S, Yang JY. Characterizing mutation-expression network relationships in multiple cancers. Comput Biol Chem. 2016 Feb 12. pii: S1476–9271

7. Lakshmanan K, Peter AP, Mohandass S, Varadharaj S, Lakshmanan U, Dharmar P. SynRio: R and Shiny based application platform for cyanobacterial genome analysis. Bioinformation. 2015 Sep 30;11(9):422–5.

8. Klambauer G, Wischenbart M, Mahr M, Unterthiner T, Mayr A, Hochreiter S. Rchemcpp: a web service for structural analoging in ChEMBL, Drugbank and the Connectivity Map. Bioinformatics. 2015 Oct 15;v31(20):3392–4.

9. Walter W, Sánchez-Cabo F, Ricote M. GOplot: an R package for visually combining expression data with functional analysis. Bioinformatics. 2015 Sep 1;31(17):2912–4.

10. Hinterberg MA, Kao DP, Bristow MR, Hunter LE, Port JD, Görg C. Peax: interactive visual analysis and exploration of complex clinical phenotype and gene expression association. Pac Symp Biocomput. 2015: 419–30.

11. Mallona I, Díez-Villanueva A, Peinado MA. Methylation plotter: a web tool for dynamic visualization of DNA methylation data. Source Code Biol Med. 2014 Jun 7;9:11. doi: 10.1186/1751-0473-9-11. eCollection 2014.

12. R Core Team. R: A Language and Environment for Statistical Computing. Vienna, Austria; 2013. Available from: http://www.r-project.org/.

13. Chang W, Cheng J, Allaire J, Xie Y, McPherson J. shiny: Web Application Framework for R; 2015. R package version 0.11.1. Available from: http://CRAN.R-project.org/package=shiny

14. Zhou T, et al. Solving the apparent diversity-accuracy dilemma of recommender systems. Proc. Natl Acad. Sci. USA 2010;107:4511–4515

15. Zhou T, et al. Bipartite network projection and personal recommendation. Phys. Rev. E Stat. Nonlin. Soft Matter Phys. 2007;76:046115.

16. Liu, Xin and Murata, Tsuyoshi. Community Detection in Large-Scale Bipartite Networks. IEEE Computer Society 2009 50–57.

17. Timothee Poisot (2015). lpbrim: Optimization of bipartite modularity using LP-BRIM (Label propagation followed by Bipartite Recursively Induced Modularity. R package version 1.0.0

18. Cheng F, et al. Prediction of drug-target interactions and drug repositioning via network-based inference. PLoS Comput. Biol. 2012;8:e1002503.

19. Alaimo S, Pulvirenti A, Giugno R, Ferro A: Drug-target interaction prediction through domain-tuned network-based inference. Bioinformatics 2013, 29(16):2004–2008.

20. Chen X, et al. Drug–target interaction prediction by random walk on the heterogeneous network. Mol. BioSyst 2012;8:1970–1978

21. Seal A, Ahn Y, Wild DJ. Optimizing drug target interaction prediction based on random walk on heterogeneous networks Journal of Cheminformatics 2015, 7:40.

22. https://aws.amazon.com/documentation/ec2/

23. David Liben-Nowell and Jon M. Kleinberg. The link prediction problem for social networks. J Comput Aided Mol Des, CIKM:556–559, 2003.

24. Mohammad Al Hasan and Mohammed J. Zaki. A survey of link prediction in social networks. Social Network Data Analytics., pages 243–275, 2011. ‘

25. David Liben-Nowell and Jon M. Kleinberg. The link prediction problem for social networks. J Comput Aided Mol Des, CIKM:556–559, 2003.

26. P. Jaccard. Etude comparative de la distribution florale dans une por-tion des alpes et de jura. Bulletin de la Societe Vaudoise des Sciences Naturelles, 37:547–579, 1901.

27. E. Ravasz, A. L. Somera, D. A. Mongru, Z. N. Oltvai, and A.-L. Barabasi. Hierarchical organization of modularity in metabolic networks. Science, 297:1553, 2002.

28. Lada A. Adamic and Eytan Adar. Friends and neighbors on the web. social networks,.Social Networks, 25(3):211–230, 2002.

29. A.L. Barabasi and R. Albert. Emergence of scaling in random networks. Science, 286:509–512, 1999.

30. Zhou T., Lu L., and Zhang Y.C. Predicting missing links via local information. Eur Phys JB,71:623–630,2010

31. Leicht E.A., Holme P., and Newman M.E.J. Vertex similarity in networks. Phys RevE, 73:026120, 2006.

32. Lu L., Jin C.H., and Zhou T. Similarity index based on local paths for link prediction of complex networks. Phys Rev E, 80:046122, 2009.

33. Katz L. A new status index derived from sociometric analysis. Psychometrika, 18:39–43, 1953.

34. Francois Fouss, Alain Pirotte, Jean-Michel Renders, and Marco Saerens. Random-walk computation of similarities between nodes of a graph with application to collaborative recommendation. IEEE Trans Knowl Data Eng, 19:355–369, 2007.

35. Sebastian Kohler, Sebastian Bauer, Denise Horn, and Peter N. Robinson. Walking the interactome for prioritization of candidate disease genes. Am. J. Hum. Genet., 82:949–958.

36. Langville A.N. and Meyer C.D. Google’s pagerank and beyond: the science of search engine rankings. Princeton University Press.

37. rCharts; [cited 4.1.2016] Available from:http://rcharts.io/

38. Barber, M. (2007) Modularity and community detection in bipartite networks. Physical Review E, 76, 066102.

39. DataTables; [cited 4.1.2016] Available from:https://www.datatables.net/

40. visNetwork; [cited 4.1.2016] Available from: http://dataknowledge.github.io/visNetwork/

41. Htmlwidgets; [cited 4.1.2016] Available from http://www.htmlwidgets.org/

42. Truchon J-F, Bayly CI (2007) Evaluating VS methods: good and bad metrics for the early recognition problem. J Chem Inf Model 47:488–508

43. Wishart DS, Knox C, Guo AC, Shrivastava S, Hassanali M, Stothard P, Chang Z, Woolsey J. DrugBank: a comprehensive resource for in silico drug discovery and exploration. Nucleic Acids Res. 2011 Jan;39(Database issue):D514-9. doi: 10.1093/nar/gkq892. Epub 2010 Oct 6.

44. Gray, Kristian A. and Yates, Bethan and Seal, Ruth L. and Wright, Mathew W. and Bruford, Elspeth A. Genenames.org: the HGNC resources in 2015.Nucleic Acids Res. 2015 Jan;43(Database issue):D1079-85. doi: 10.1093/nar/gku1071. Epub 2014 Oct 31.

45. https://www.sqlite.org/

46. Durinck S, Moreau Y, Kasprzyk A, Davis S, De Moor B, Brazma A and Huber W (2005). BioMart and Bioconductor: a powerful link between biological databases and microarray data analysis. Bioinformatics, 21, pp. 3439–3440.

47. Yu G, Wang L, Han Y and He Q (2012). clusterProfiler: an R package for comparing biological themes among gene clusters. OMICS: A Journal of Integrative Biology, 16(5), pp. 284–287

48. Yu G and He Q (2016). ReactomePA: an R/Bioconductor package for reactome pathway analysis and visualization. Mol.BioSyst., 2016,12, 477–479

49. Yamanishi Y., Araki M., Gutteridge A., Honda W., Kanehisa M.. Prediction of drug–target interaction networks from the integration of chemical and genomic spaces. Bioinformatics 2008;24:i232–i240.

50. Devtools by Hadley Wickham. https://github.com/hadley/devtools

51. RDKit: Cheminformatics and Machine Learning Software. 2013,: -[http://www.rdkit.org], [http://www.rdkit.org]

52. Bleakley K, Yamanishi Y (2009) Supervised prediction of drug–target interactions using bipartite local models. Bioinformatics 25:2397–2403

